# A detailed protocol for generating spike trans-complemented SARS-CoV-2 replicons

**DOI:** 10.1101/2024.07.28.605443

**Authors:** Qi Yang, Balaji Manicassamy, Sriram Neelamegham

## Abstract

Multiple approaches have been implemented for basic science studies that attempt to investigate SARS-CoV-2 biology or virology in biosafety level-2 setting. These include pseudotyped-virus based on lentivirus and vesicular stomatitis virus, virus-like particles that only contain the SARS-CoV-2 structural proteins, and single-cycle replicons. Among these, the single-cycle replicons most closely resemble the authentic virus as they essentially include the full viral genome, except for essential elements required for active virus multiplication. In this regard, we previously developed a SARS-CoV-2 replicon system where a Gaussia Dura luciferase-P2A-mNeonGreen reporter cassette replaced viral spike. In this short paper, we present an optimized protocol for the use of this reagent that overcomes previous technical limitations. We demonstrate that co-transfection of this bacmid along with spike plasmid, using the improved protocol, yields high-quality spike bearing SARS-CoV-2 virus particles with single-cycle infectivity. Due to the nature of bacmid construction, this approach is particularly useful for studying the impact of spike mutagenesis on virus evolution in BSL-2 setting.

## MAIN TEXT

Studies of SARS-CoV-2 require BSL3 biosafety for handling due to high pathogenicity and transmissibility, which hampers basic science investigations. To enable such study in BSL2 setting, pseudovirus and virus-like-particles have been developed (*1, 2*). These solutions, however, only partially mimic the original virus due to ectopic budding and lack of complete viral genome. As an alternative, single-cycle infectious SARS-CoV-2 virus replicon particles (VRPs) were developed.

These methods exclude essential viral genome components needed to assemble infective virus. Examples of such components include non-structural protein 15 (nsp15) (*3*), spike and envelope together with membrane (*4*), ORF3 and envelope (*5*), nucleocapsid (*6*), spike alone (*7, 8*) or the breakdown of SARS-CoV-2 genome into multiple plasmids (*9*). Upon trans-completing the missing essential components with the base constructs, virions are produced that result in single-cycle infection without producing infective virions. While valuable, producing such particles can be methodologically complicated due to the need to perform *in vitro* transcription (*8*), use of specialized cells (*5*), stoichiometric addition of multiple plasmids (*9*) or other considerations explained below.

With the goal of creating systems that allow studies of spike glycosylation (*2, 10*), we previously developed a SARS-CoV-2 bacmid replicon where a Gaussia Dura luciferase-P2A-mNeonGreen reporter cassette replaced viral spike (*7*). These ΔS virus-replicon-particles (ΔS-VRPs) when trans-complemented with viral glycoproteins resulted in single-cycle viral particles. While previous studies developed VSV-G pseudotyped VRPs, incorporation of spike only yielded virions with low infectivity. We have overcome this limitation recently and developed a straightforward protocol that is described in this submission (details in **Supplemental Material**).

Several approaches were attempted to produce VRPs trans-complemented with spike.

Among them, in some studies, we produced ΔS-VRPs with VSV-G (*7*), and then co-infected cells with this stock virus along with SARS-CoV-2 spike plasmid to attempt to generate ΔS-VRPs bearing spike. This was attempted both using cells lacking human ACE2 (i.e. wild-type HEK293T without Angiotensin-converting enzyme 2) and cells bearing ACE2 (Vero E6, A549-ACE2-TMPRSS2, 293T-ACE2). Paradoxically, none of these produced ΔS-VRPs with spike since viral transmission was low in the absence of ACE2 in producer cells, while ACE2 expression led to extensive syncytia formation and death resulting in low viral yield. Various protocols were then attempted to co-transfect replicon and spike plasmids into the producer cells. The optimized conditions used 50% hepatocyte-derived carcinoma Huh7.0 cells and 50% human embryonic kidney 293T cells as outlined in Supplemental Methods (**Fig. 1A**). ΔS-VRP SARS-CoV-2 spike viral particles generated in this manner resulted in host cell infection as assayed using microscopy measurement of mNeonGreen positive cells **(Fig. 1B)** and Gaussia luciferase luminescence (**Fig. 1C**). Viral entry was spike-dependent as VRPs expressing spike presented >10-fold higher signal, compared to ΔS-VRP lacking spike (**Fig. 1C)**. Unlike spike VRPs, VSV-G VRPs exhibited higher transmission and thus higher viral titers. Overall, the letter presents a simple and detailed protocol for generating ΔS-VRPs complemented with spike protein. As part of ongoing efforts, this approach is being used to study the role of glycosylation on spike from SARS-CoV-2 and also other *Coronavirdae* family members.

**Figure 1.**
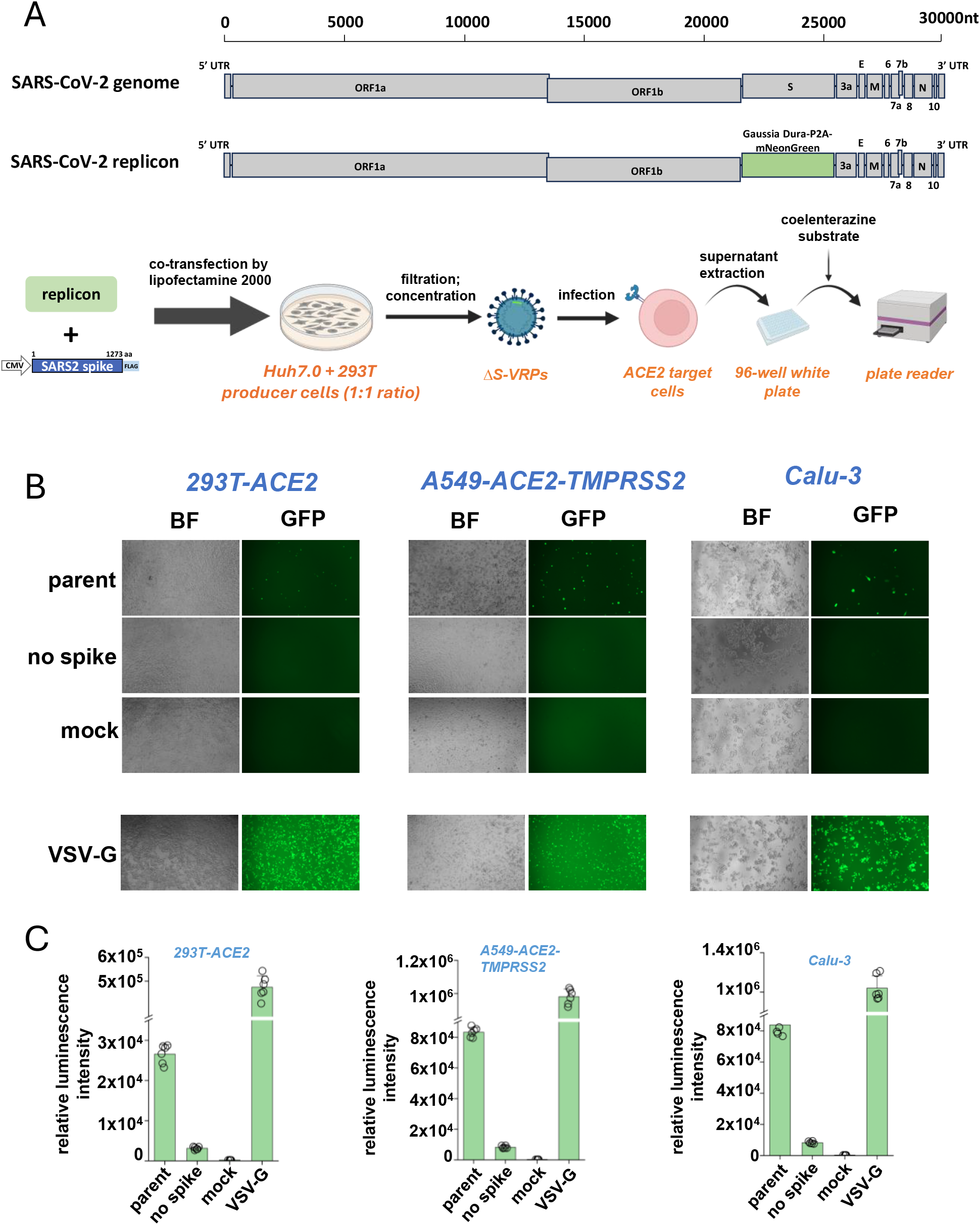
**A**. Method for producing ΔS-VRP bearing SARS-CoV-2 spike. **B**. mNeonGreen signal upon transduction of various cells bearing human ACE2. **C**. Luminescence measurements using supernatant of the same samples from **B**.

