## Supplemental Methods for "A detailed protocol for generating spike trans-complemented SARS-CoV-2 replicons"

### SUPPLEMENTARY PROTOCOL

Bring all reagents described below to room temperature before use.

### MATERIAL

**Transfection reagent**: lipofectamine 2000 (Thermofisher # 11668027)

**E. coli**: DH10B bacteria (New England Biolabs # C3019I)

**Producer cells**: hepatocyte-derived carcinoma Huh7.0 cell (ATCC), human embryonic kidney 293T cells (Takara bio)

**DNA**: SARS-CoV-2 replicon bacmid (*1*), SARS-CoV-2 parent spike (*2*), VSV-G glycoprotein (Addgene: 12259)

**Protein concentrator**: 100kDa polyethersulfone (PES) protein concentrator (Fisher Scientific # 88533)

**Buffer**: HEPES buffer (30mM HEPES, 110mM NaCl, 10mM KCl, 2mM $MgCl_2 \cdot 6H2O$, 10mM glucose, pH 7.3)

**Luciferase assay kit**: Gaussia Luciferase Glow Assay Kit (FisherScientific # 16160)

### METHODS

#### A. Scaling up bacmid DNA

1. Thaw the DH10B bacteria on ice and add 25 µl of bacteria to 1.5 ml tube.
2. Add 50 ng of bacmid to the bacteria and incubate for 30 min on ice.
3. Heat shock at 42°C for 30s. Immediately keep on ice for recovery for 5 min.
4. Add 1 ml of SOC media.
5. Incubate at 33°C for 2 hours.
6. Plate 100µl of bacteria in agar plate with chloramphenicol (25 µg/ml) and incubate for 16-20 hours.
7. Pick colonies and do a 1 ml starter culture in 2xYT media/chloramphenicol (25 µg/ml).
8. Incubate at 33°C for 2 hours in a shaker.

9. Use the start culture to inoculate 450-500 ml maxiprep culture. Incubate for 18-20 h and proceed with maxiprep for scaleup using NucleoBond Xtra Maxi kit.

**B. ∆S-VRP production and spike trans-complementation**

1. Mix 12 µg replicon bacmid DNA and 4 µg spike or VSV-G plasmid in 500 µl serum reduced OPTI-MEM (Invitrogen).
2. Mix 48 µl lipofectamine 2000 in 500 µl serum reduced OPTI-MEM.
3. Add diluted replicon bacmid/spike or VSV-G DNA into lipofectamine 2000 vial and mix thoroughly. Incubate at room temperature for 15-20 min.
4. Mix $4\times10^6$ hepatocyte-derived carcinoma Huh7.0 cells and $4\times10^6$ human embryonic kidney 293T (HEK293T) and plate in 100 mm petri dish containing 15 ml DMEM (ThermoFisher # 11995-073) with 10% FBS (Fetal bovine serum).
5. Add DNA/lipofectamine 2000 mix to cells dropwise with gentle rocking.
6. 6-8 hours post transfection, switch media to 15 ml DMEM containing 2% FBS. Wait 72 h to allow viral production.
7. Collect supernatant, pellet any cells using 4000g for 5 min, and filtered using 0.45 µm PES filters to remove cell debris.
8. Add clarified supernatant to 100 kDa protein concentrator, buffer exchanged against HEPES buffer to deplete residual luciferase signal from producer cells and concentrate by 30-fold to a final volume of 500 µl. The ∆S-VRPs can be directly used for viral entry assay or for long-term storage at -80°C.

**C. ∆S-VRP infection studies**

1. Suspend target/host cells at $10^7$ /mL DMEM containing 2% FBS. Add 50 µl ∆S-VRPs, 8 µl cells and 6 µl polybrene (8mg/ml stock).
2. Mix these components in 1.5 mL Eppendorf tube at room temperature for 30 min, and then transfer into 96-well plates with additional 50 µl DMEM containing 2% FBS.

3. After overnight incubation, perform fluorescence investigation of target cells, and collect supernatant for Gaussia luciferase assay.

**D. Gaussia luciferase assay**

1. Gaussia luciferase working solution is prepared by following the manufacturer's guidance.
2. Pipette 50 µl supernatant from the target cells and add to a white 96-well plate with round bottom. 50 µl working solution is then added into the supernatant.
3. Samples are incubated for 10 min at room temperature for stabilizing the signal.
4. Measure luminescence using plate reader.
